# Learning structural motif representations for efficient protein structure search

**DOI:** 10.1101/137828

**Authors:** Yang Liu, Qing Ye, Liwei Wang, Jian Peng

## Abstract

**Motivation:** Understanding the relationship between protein structure and function is a fundamental problem in protein science. Given a protein of unknown function, fast identification of similar protein structures from the Protein Data Bank (PDB) is a critical step for inferring its biological function. Such structural neighbors can provide evolutionary insights into protein conformation, interfaces and binding sites that are not detectable from sequence similarity. However, the computational cost of performing pairwise structural alignment against all structures in PDB is prohibitively expensive. Alignment-free approaches have been introduced to enable fast but coarse comparisons by representing each protein as a vector of structure features or fingerprints and only computing similarity between vectors. As a notable example, FragBag represents each protein by a “bag of fragments”, which is a vector of frequencies of contiguous short backbone fragments from a predetermined library.

**Results:** Here we present a new approach to learning effective structural motif presentations using deep learning. We develop DeepFold, a deep convolutional neural network model to extract structural motif features of a protein structure. Similar to FragBag, DeepFold represents each protein structure or fold using a vector of learned structural motif features. We demonstrate that DeepFold substantially outperforms FragBag on protein structural search on a non-redundant protein structure database and a set of newly released structures. Remarkably, DeepFold not only extracts meaningful backbone segments but also finds important long-range interacting motifs for structural comparison. We expect that DeepFold will provide new insights into the evolution and hierarchical organization of protein structural motifs.

**Availability:** https://github.com/largelymfs/DeepFold

## 1 Introduction

The comparison of protein structures has been a fundamental and widely applicable task in structural biology. Given a new protein with unknown functions, searching similar protein structures from existing databases is a critical step for predicting its function. It is particularly valuable when the query protein shares little sequence similarity with existing ones, where sequence alignment algorithms, such as BLAST[4], cannot easily identify evolutionary relationships. Protein structural search against a database is often prohibitively expensive because performing pairwise structure alignments against all proteins in a database is computationally linear to the number of protein structures in the database, which is usually large. Due to the advances in crystallography and cryo-EM, the Protein Data Bank[1], a database of solved protein structures, has recently been increasing rapidly, with new protein structures solved almost every day, though the number of possible protein folds is thought to be bounded. Structural classifications of PDB, including SCOP[7] and CATH[18], enable us to explore the hierarchical organization of protein structure space and provide useful guidelines on the relationship between protein structure and functions by identifying structural neighbors. However, the classification of structural neighbors may not be optimal, as there are neighbors, though functionally and structurally similar, classified differently. Thus, fast and accurate structural search against a large-scale protein structure database is still a challenge.

Arguably the most popular structural comparisons approaches are structural alignment-based methods [30, 13, 16, 24, 22]. Pairwise structural alignment algorithms take 3D coordinates of two protein structures and optimize a predefined geometric similarity measure using heuristics, such as simulated annealing, dynamic programming and local search. Although these approaches can provide residue-resolution alignments with high accuracy, they are often computationally expensive. Such computational cost makes large-scale structural search not practical when coping with a very large database since the runtime of such structural alignment algorithms scales linearly to the number of proteins in the structure database.

Another entirely different approach for structural search is to represent protein structures using structural features as 1D vectors and then perform similarity calculation of such vectors without performing an alignment. The idea of alignment-free algorithms was initially introduced for protein sequence comparisons. A feature vector is computed to represent each protein sequence for fast cosine- or Euclidean distance-based similarity calculations. For instance, CD-HIT computes a histogram of k-mers for fast sequence clustering [9]. Other examples include k-mer mismatch kernels for homology search and compositional methods for metagenomic binning [6, 10]. In many applications, these methods can provide fast comparisons with comparable performance to the alignment-based methods. Recently, several alignment-free protein structural comparisons have been proposed [3, 14, 31]. By representing each protein as a vector of structure features or fingerprints, we can compute the similarity between vectors to find structural neighbors in a large database. As a notable example, FragBag represents each protein by a “bag of fragments”, which is a vector of frequencies of a set of predefined contiguous short backbone fragments [3]. Although it achieves comparable accuracy to some structural alignment algorithms, the performance of FragBag is not satisfactory when the database becomes large. A possible reason is that FragBag only considers contiguous backbone fragments, which captures only local property of the whole structure. This limitation is sub-optimal because important long-range interacting patterns are ignored.

Here we present a new approach to learning an innovative structural motif presentation using deep learning. Inspired by FragBag, we hope to further generalize it by learning conserved structural motif for protein structure representation. We develop DeepFold, a deep convolutional neural network model to extract structural motif features of a protein structure from its *C*_*α*_ pairwise distance matrix. This neural network model extracts local patterns from residue contact patterns and composes low-level patterns into high-level motifs from layer to layer. From this neural network, we can represent each protein structure/fold using a vector of structural motifs and perform the structural search by only computing vector similarity. This neural network model is trained in a supervised manner by fitting TM-scores [29, 30], a structural similarity score, between existing protein structures from a nonredundant SCOP database. We demonstrate that DeepFold substantially outperforms three existing alignment-free methods, FragBag [3], SGM [14] and SSEF [31], on protein structure search for a set of newly released PDB structures in 2016. DeepFold achieves improved structural search accuracy and obtains better structural neighbors. We also show that DeepFold not only extracts conserved backbone segments but also identifies important long-range interacting structural motifs in the representation. We expect that DeepFold will provide new insights into the evolution and hierarchical organization of protein structural motifs.

## 2 Background: protein structure comparison and retrieval

### 2.1 Structural alignment

Pairwise structural alignment algorithms take two protein structures as input and identify geometrically aligned substructures according to a predefined structural similarity score. Notable examples of structural alignment algorithms include TM-align [30], Combinatorial Extension (CE) [16], DeepAlign [22], Matt [13] and FATCAT [24]. It is worth noting that, given a structural similarity score, sophisticated optimization and sampling techniques are usually required to optimize the score and find the best alignment and substructures. As a result, the computation of aligning two structures is expensive. Therefore, using structural alignment algorithms for structural neighbor retrieval would require *O*(*n*) pairwise structural alignments, where *n* is the size of the structure database used for retrieval. By February 1st of 2017, there have been more than 117, 000 protein structures deposited in the PDB. Even the non-redundant SCOP and CATH include more than 13, 000 structural domains. Thus, naively applying structural alignment algorithms for protein structure retrieval is prohibitively expensive.

### 2.2 Alignment-free methods

In contrast to structural alignment algorithms, alignment-free methods have been proposed for accelerating structural neighbor search. Instead of optimizing a complex score for identifying geometrically similar substructures, alignment-free methods represent each protein as a set of structural patterns or fingerprints encoded in a feature vector and compare these representations via very simple similarity calculations like cosine- or Euclidean similarity. For example, Zotenko et al. [31] count the frequencies of 1, 500 secondary structure triplets (SSEF) as a vector for representing a protein structure. Scaled Gaussian Metric (SGM) [14] represent a protein structure using 30 global backbone topological measures. Recently, Kolodny and co-workers developed FragBag [3], a ”bag-of-words” vector representation including frequencies of local contiguous fragments in the protein backbone. The optimal performance was achieved with a library of 400 backbone segments, each with 11 contiguous residues. Although FragBag shows improved retrieval performance over previously existing alignment-free methods, the accuracy of FragBag is still not satisfactory compared to advanced alignment-based methods. Possible reasons are that 1) the backbone fragment library may not be optimal and that 2) the long-range interacting patterns, which are known to be highly important in discriminating different protein fold, were not considered.

## 3 Learning structural motifs for protein structure representation

We propose DeepFold, a novel deep learning based method to project tertiary protein structures into a low-dimensional vector space. Different from conventional protein structure filtering methods like FragBag, our method does not need a predefined fragment library but automatically identifies relevant local structural protein motifs and long-range interacting motifs. Benefiting from the powerful representation learning ability of deep neural network, DeepFold is able to construct a more powerful protein structure representation, achieving a better performance on the task of protein structural searching.

### 3.1 Protein structure comparison

Suppose we have a query structure *x*_*Q*_ and two candidate template protein structures *x*_*A*_, *x*_*B*_. By applying the DeepFold neural network on these structures as a feature extractor, we obtain three corresponding structural fingerprint vectors **v**_*Q*_, **v**_*A*_, **v**_*B*_. With these fingerprints, we compute the similarity of each candidate to the target query by taking the cosine similarity of their corresponding vectors and ranking all candidates from a structure database accordingly. A schematic diagram of this procedure can be found in Figure 1. Thus, how to effectively represent each protein into a low-dimensional vector space is quite critical for enabling accurate protein structure search.

**Figure 1:**
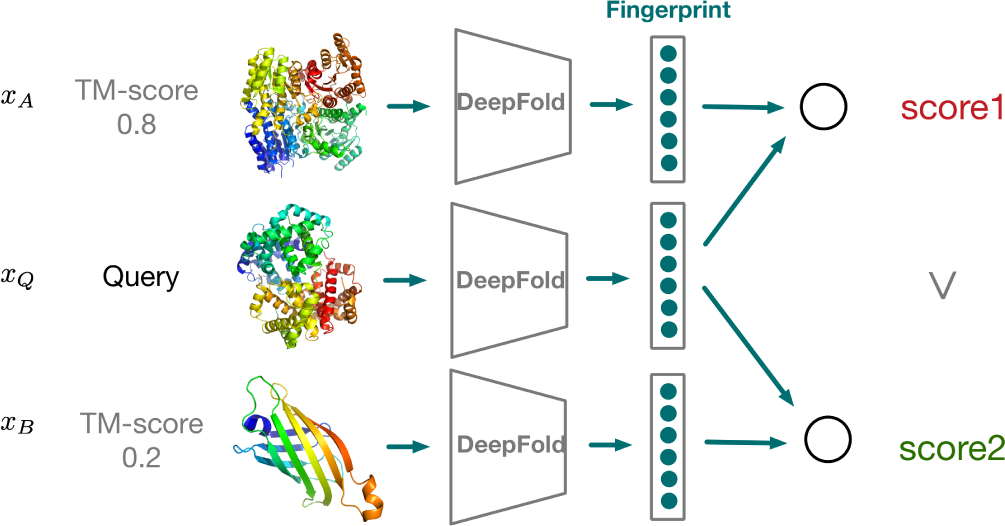
Alignment-free structure comparison. The query protein structure is mapped into a low-dimensional fingerprint vector. We then compute a cosine similarity score between the query fingerprint vector and the vector of a template structure. Similar structures share high similarity scores, while dissimilar structures share low similarity scores.

### 3.2 Deep convolutional neural network

Deep convolutional neural networks (DCNNs) are natural for learning hierarchical representations of image data. Multiple layers of convolutional filters are constructed for identifying local patterns in a nested manner. Inspired by the recent successes of DCNNs in computer vision and image processing, we propose to develop a deep convolutional neural network model, DeepFold, to learn structural motifs for structural comparison.

Given a tertiary structure of a protein, we first calculate the *C*_*α*_ pairwise distance matrix as the raw representation feature, aiming at preserving the geometric information of the input protein structure. We denote the proteins structure as *x* and its pairwise distance matrix as **D**, each entry **D**_*ij*_ being the *C*_*α*_ distance between *i*^*th*^ residue and *j*^*th*^ residue. Due to the existence of missing residues in protein structures, directly taking the distance matrix as input may later cause numerical issues in the neural network. So instead, we use the following tensor as input.

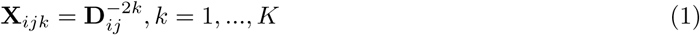

where **X**∈**R**^*L*×*L*×*K*^ and *K* is an integer indicating the inverse power of the squared distances. It is worth noting that the choice of this inverse power series of distances is similar to several distance-dependent approximations in force field energy functions, such as van de Waals and electrostatic energies [23, 2].

**Figure 2:**
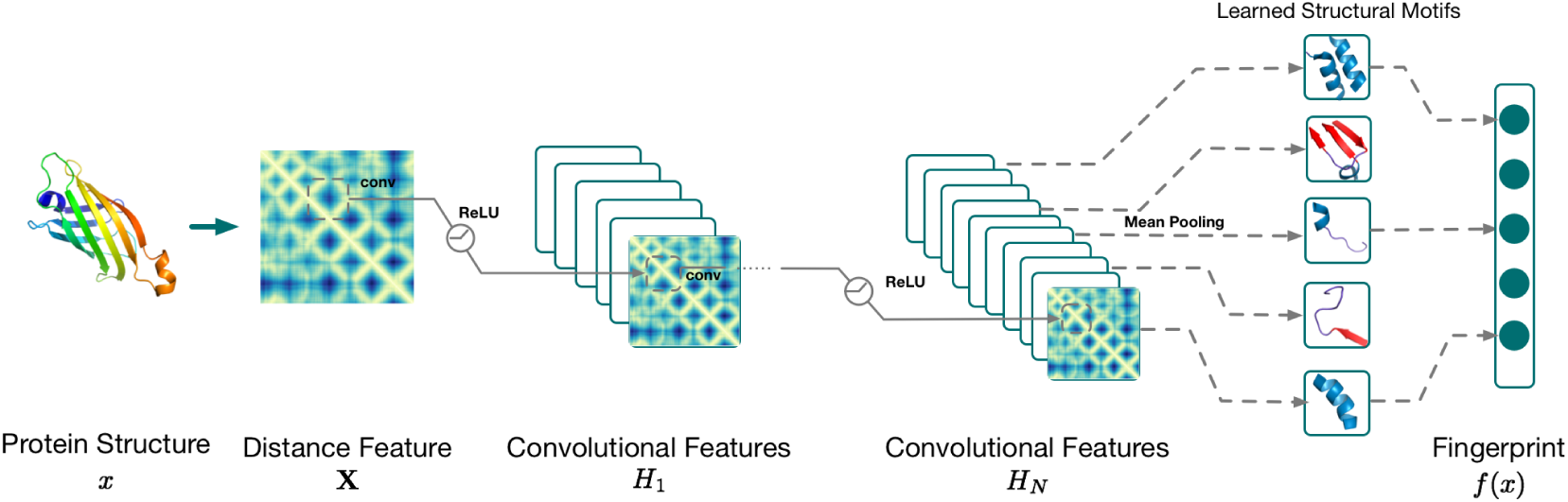
Deep neural network structure of DeepFold: Given a protein structure, we firstly compute the raw distance features. Then we feed the features into a deep convolutional neural network that consists of N convolutional blocks. Inside each block, we apply a convolutional layer, followed by a ReLU nonlinear layer. The final output of DeepFold is a vector representation of structural motifs.

As shown in Figure (2), a deep neural network, DeepFold, takes the transformed input feature of **X**, followed by a sequence of blocks of transformations. Each block contains the stacked layers including a convolutional layer and a nonlinear transformation ReLU layer. After the last convolutional layer, we apply a mean-pooling layer and L2 normalization to aggregate features and obtain the final fingerprint representation *f* (**X**) of the input protein structure. Taken as a whole, our deep net is a non-linear mapping from a high-dimensional structural input **X** to a low-dimensional fingerprint representation *f* (**X**). Specifically, a convolutional layer takes the output from the previous layer *H*_*n–*1_∈ **R**^*L*_*n–*1_×*L*_*n*–1_×*K*_*n*–1_^ as input and compute output *H*_*n*_∈ **R**^*L*_*n*_×*L*_*n*_×*K*_*n*_^ in the following way. For each *k* ∈ {1, 2,‥, *K*_*n*_}

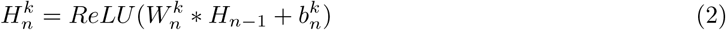

where for the n-th convolutional layer, 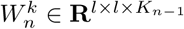 is a weight tensor of *K*_*n–*1_ convolutional filter weights of size *l* × *l*. The function operator (*) is 2-dimensional convolution operator over the first two dimensions of the input *H*_*n–*1_, and *b*_*n*_ is a bias term. Intuitively, a filter in the first convolutional layer can be seen as a local feature extractor on the inverse pairwise distance matrix with the goal to capture important local residue contact patterns as a structural motif. In higher layers, a filter can be seen to compose low-level structural motifs into high-level motifs. However, note that the convolutional operation essentially performs dot products between the filters and local regions of the input features. A common implementation of the convolutional layer is to take advantage of this fact and formulate it as matrix multiplications. The output dimensions *L*_*n*_ × *L*_*n*_ depend on the stride which further reduces the feature dimensionality by controlling how the convolutional filters slide across the input of each layer. After each convolutional layer, a Rectified Linear Unit (ReLU) function [8] is applied for nonlinear activation. The ReLU can be implemented by simply thresholding a matrix of activations at zero by *ReLU* (*h*) = max(0*, h*). Given any tensor/matrix/vector input *h*, ReLU takes elementwise max operations. Compared to traditional logistic or tanh functions, ReLU is found to greatly accelerate the convergence of stochastic gradient descent due to its linear, non-saturating form.

After the last convolutional layer, we get *K*_*N*_ feature maps with the size of *L*_*N*_ × *L*_*N*_. Then for each feature map *i* ∈ {1, 2,‥, *K*_*N*_}, we extract the diagonal elements of each feature map and take the mean pooling (that is, calculating the mean value of diagonal elements). Therefore, we get the *K*_*N*_ -dimensional vector representation of *f* (*x*). For optimization efficiency, each vector is normalized with L2 Norm so that all the vectors are projected on a sphere || *f* (*x*)|| _2_ = 1. Similar to the FragBag representation, DeepFold maps an input tertiary structure into a low-dimensional one-dimensional fingerprint vector but this representation is parameterized via a deep neural network, with each dimension encoding a specific structural motif. Therefore, DeepFold is more powerful than FragBag in that it extracts not only contiguous backbone fragments but also the long-range interacting motifs, which are completely omitted by FragBag. One can easily see that FragBag becomes a special case of DeepFold if we only consider the near-diagonal region of the pairwise distance matrix as input. Our in-house experiments showed that the learning convergence of this special case is poor in practice.

### 3.3 Learning to compare protein structures

Given two proteins *x*_*A*_ and *x*_*B*_, with their normalized representations *f* (*x*_*A*_) and *f* (*x*_*B*_), we expect the similarity between *f* (*x*_*A*_) and *f* (*x*_*B*_) can reflect the structural similarity between *x*_*A*_ and *x*_*B*_. So if two proteins have very similar structures, we should map their features *f* (*x*_*A*_) and *f* (*x*_*B*_) to be close to each other, while if *x*_*A*_ and *x*_*B*_ have different structures, we should push their fingerprint features far from each other. Although pairwise structural alignments are slow in protein structural retrieval, we can precompute them on a training dataset and use the resultant structural similarity scores to guide the training of DeepFold. So our key idea here is to fit structural comparison scores using powerful deep neural networks, thus enabling fast alignment-free comparison and retrieval for new query proteins. To implement this idea for training DeepFold, we use the cosine similarity score of *f* (*x*_*A*_) and *f* (*x*_*B*_) and apply the well-known max-margin ranking loss function [21] to discriminate structurally similar proteins from dissimilar ones.

To apply the max-margin rank loss function, we firstly define what positive (similar) and negative (dissimilar) pairs of proteins are. In this work, we use the structural alignment program TM-align [29] to measure the structural similarity of two proteins. Based upon a dynamic programming algorithm, TM-align performs structural alignment of the two input structures to produce an optimal superposition and return a TM-score that scales the structural similarity in the range of [0, 1]. Therefore, for a protein *x*_*A*_, we define all pairs (*x*_*A*_, *x*_*B*_) from the database with TM-score higher than *p* T M_max_*(*x*_*A*_) as positive pairs, where *T M_max_*(*x*_*A*_) is the maximal TM-score between *x*_*A*_ and other proteins in the database, and *p* is a hyper-parameter chosen to be 0.9. So only very similar structures in the database are considered to construct positive pairs in training. For all other pairs that have scores smaller than this threshold, we consider them as negative pairs. It is important to note that other structural similarity scores or structural alignment algorithms can be used here. We choose TM-align/score as it has been shown to be both fast and accurate in structural classification and often more robust than many other structural similarity scores [30].

With the defined positive and negative pairs, our target is to make the margin between all positive pairs and negative pairs as large as possible. Formally, for a specific protein *x*_*A*_, suppose the set of all positive pairs is 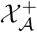 while the negative set is 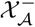. Thus, the margin loss could be defined as :

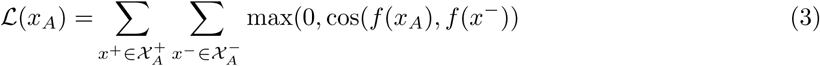

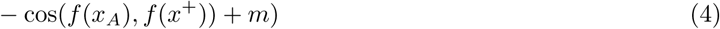

where *m* is a small positive margin value, which is chosen as 0.1 in this work. Then, the objective is to minimize this margin loss and learn the parameters {*W, b*} in all layers of the neural network as defined in the above.

#### Efficient online training

In above Equation 4, to optimize the loss function, we need to sum over all positive and negative training pairs and minimize the rank loss function. However, the number of training pairs can be very huge, and thus it is not feasible to optimize them all at one time. In addition, the imbalance of positive and negative pairs pose another challenge in training. Here, we use the stochastic gradient descent to only randomly take a small batch of training samples at in each iteration. Specifically, we use mini-batches to do the feed-forward passes and back-propagation in each iteration. Motivated by [15], we design an simple yet effective sampling method to obtain a fast empirical convergence by selecting effective pairs. Given a protein *x*_*A*_, we consider the most effective positive pairs and negative pairs 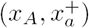 and 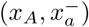 accordingly as:

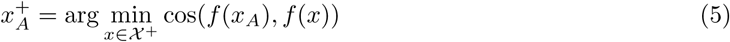

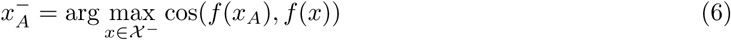

Intuitively, we identify those positive pairs that are easier to get confused with negative pairs (furthest positive pairs) and those negative pairs that are easier to get confused to positive pairs (nearest negative pairs) within each mini-batch. If mini-batches may not have any positive pair, we add at least one positive pair for each protein in the mini-batch to balance the training. Also, inside each mini-batch, we identify negative targets that are most similar to the query and incorporate the margin between all positive pairs and these k negative pairs to make the training process more effective.

### 3.4 Implementation details

The first convolutional layer consists of 128 filters of size 12 × 12 with a stride 2 × 2. After that, we connect it with the second layer of 256 kernels of size 4 × 4 with a stride of 2 × 2. From the third layer, we stack 3 identical convolutional layers, each with the size of 4 × 4 and a stride 2 × 2. Finally, the output of stacked layers is linked to 400 filters with a size of 4 × 4 and a stride of 2 × 2. The total dimension of 400 is selected to match the size of the vector representation used in FragBag. Based on the features extracted, we capture only diagonal elements and perform mean pooling for each filter. Then we conduct L2 normalization and project all final fingerprints into the space of sphere ||*x* ||_2_ = 1.

DeepFold is implemented with the Python library Lasagne^1^ based on Theano platform[20]. In the optimization, we apply stochastic gradient descent(SGD) with a momentum of 0.9. The learning rate decay scheme is chosen with AdaGrad[5] algorithm. The mini-batch size is chosen as 64 considering the memory usage on the graphics card. Within each mini-batch, we sample one positive instance for each protein and compute top 10 hardest negative instances in the margin loss inside each mini-batch. To make the model more robust, we used Dropout[19] after each ReLU layer, which could be viewed as an ensemble trick to enhance the generalization ability of the model. During training, we monitor the retrieval accuracy using the validation set as queries and the training set as the database. Totally, we train each model with 100 epochs and select the one with best evaluation accuracy. Then we report the performance tested with testing fold as queries and all of the training data as database on this selected model. All the experiments are performed on a workstation with 256GB RAM and a NVIDIA Titan X graphics card with 12GB memory.

## 4 Results

### 4.1 Comparison to existing alignment-free structural retrieval methods

#### Experimental settings

We construct a sequence-nonredundant structure dataset by filtering out protein structures with 40% sequence identity in the latest SCOP database (version 2.06) [7]. The filtered database includes 13, 546 representative protein domains, indexed by the manually curated SCOP taxonomy classification.

We compare our proposed DeepFold with FragBag [3], the existing state-of-the-art alignment-free method. With the goal of making a fair comparison, we set the dimensionality of DeepFold representation to be 400, which is the same parameter used in the default setting of FragBag. Also, we also compare our approach with two other previous alignment-free methods, including SGM [14] and SSEF [31], which

were introduced much earlier. Furthermore, to test whether the long-range patterns are helpful, we also build a simplified local DeepFold model as a baseline, which only extracts fragmental features from consecutive residuals, the same as FragBag. This method is denoted by DeepFold(L).

For assessment, we pick structures with TM-Scores that are no less than 0.9 of the highest TM-Score achievable by structures in the database as the True structural neighbors to a target structure. It is natural to train and/or test DeepFold with other structural similarity metrics such as GDT TS[27][26], MaxSub[17] and lDDT[12].To evaluate the accuracy of structural retrieval, we compute the Receiver Operating Characteristic(ROC) curve for each method and calculate the average area under ROC curve (AUROC) of all query proteins in the validation set. A more effective algorithm should have a larger AUROC. In addition to the ROC curve, we also plot the hit@top-*k* curve, which has been widely used for evaluating ranking performance in information retrieval [28]. The hit@top-*K* is calculated as the percentage of queries in which at least one positive template (i.e. TMscore *>* 0.9\**T M_max_*(*query*)) appear in the top-*K* retrieved list from the dataset. We choose the top-*K* hit rate as the metric motivated by the fact that if there is a very similar structural neighbor ranked within top *K*, we can apply a structural alignment algorithm (such as TM-align[30]) to identify it with at most *K* pairwise alignments. Besides the curves mentioned above, we also report the area under the precision-recall curve(AUPRC) along with hit@top-1,hit@top-5,hit@top-10 in the table, providing alternative metrics for comparison. In addition, we also compute hit@top-K accuracy by the SCOP classification, i.e. whether two structures are within the same family classification.

#### Results on SCOP data

Since DeepFold is trained by supervised learning, in contrast to other unsupervised alignment-free structural comparison methods, we firstly design a rigorous cross-validation scheme to evaluate the performance on the SCOP structure database more robustly. Specifically, we randomly split all structures from the SCOP database into 5 subsets and perform a 5-fold cross validation. In each fold, we only use pairwise TM-scoresbetween proteins from the training set as ground truth to train DeepFold model, and for evaluation, we utilize each protein in the validation set as the query to search in the training set and report the retrieval accuracy. We show ROC curves and Hit curves on cross-validation and the comparison results in Fig. 3, Fig. 4 and Table 1 respectively.

**Figure 3:**
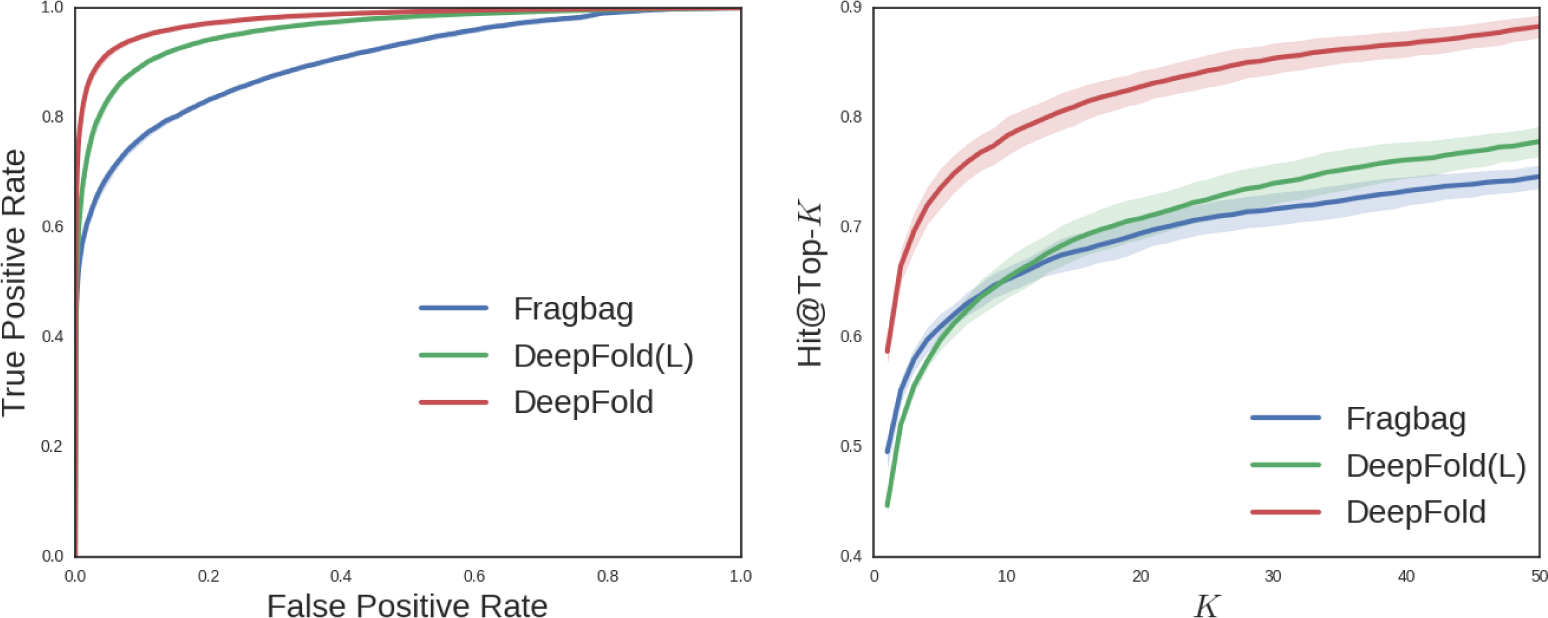
ROC curves and Hit@Top-*k* curves on SCOP data. The performance of DeepFold is compared with FragBag and a simplified version of DeepFold with only local backbone segmental features.

**Figure 4:**
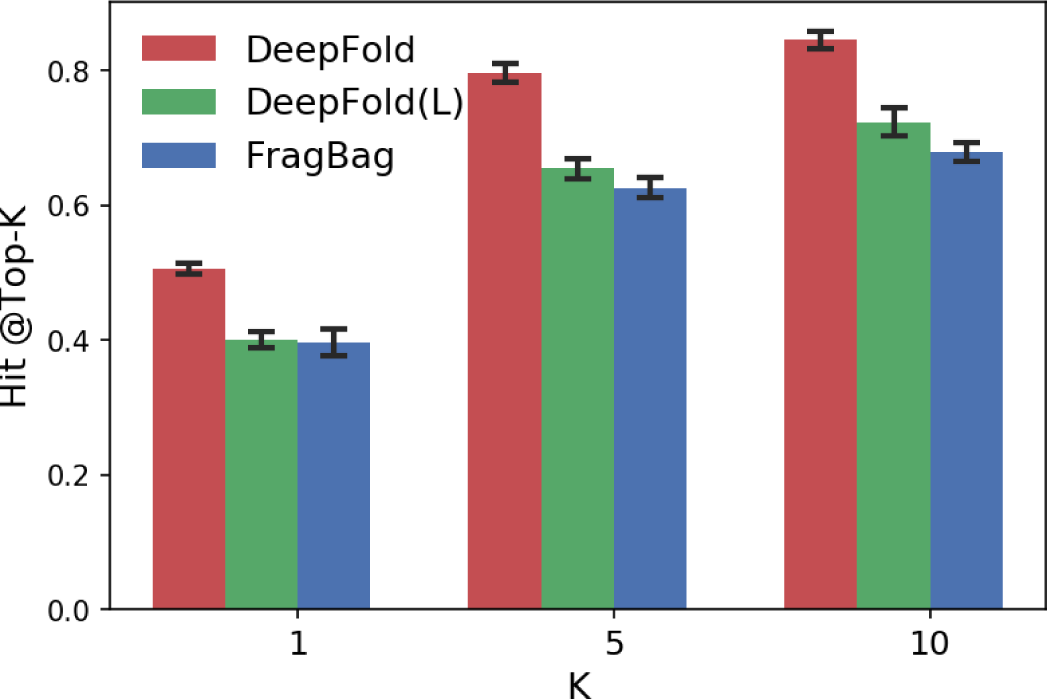
Hit@Top-*k* according to SCOP family classification. The performance of DeepFold is compared with FragBag and DeepFold(L).

**Table 1:**
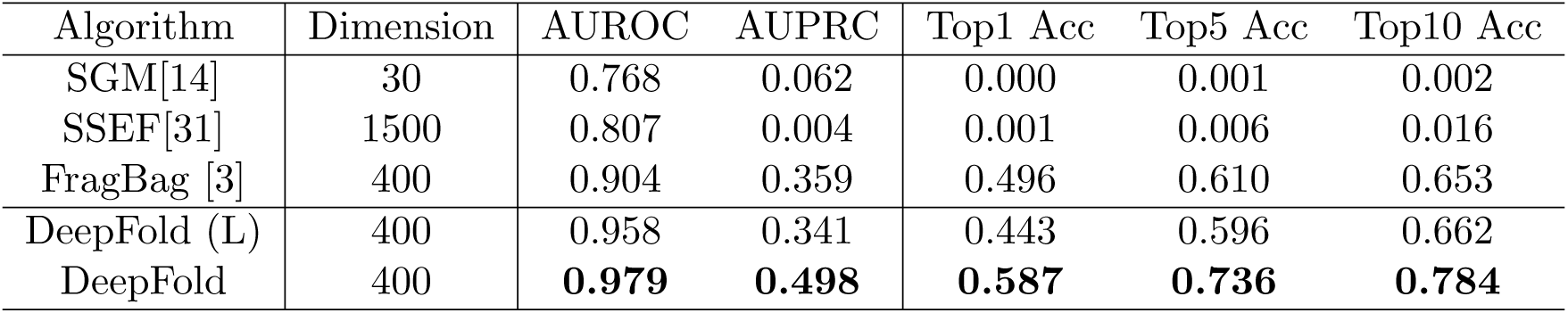
Performance comparison on the SCOP data.

According to the average ROC Curve of these algorithms, our method significantly outperforms other alignment-free methods. Our algorithm achieves a much higher true positive rate at the same level of the false positive rate. Also, our algorithm has a higher AUPRC score, indicating DeepFold is better at ranking similar structures on the top. Moreover, according to the top-1, top-5, top-10 accuracy, our algorithm has remarkable improvement compared with the strongest baseline, FragBag. From Fig 4, we also observe the significant improvement in the accuracies determined by the SCOP family classification, indicating our method is consistantly better and not overfitted to the metric used for training. The improvement we observed is probably because our algorithm could automatically learn structural motif which is hard to be computed by human clustering or there maybe consist of several unknown protein structures which have not be explored by the human before. Furthermore, FragBag treats each structural motif equally which is not consistent with biological insights. Nevertheless, our DeepFold can learn the weight for each filter automatically so that each motif is attached with a weight. In addition, our DeepFold(L) outperforms FragBag according to AUROC but slightly worse according to hit@top-k and AUPRC. We think the possible reason is that Fragbag is constructed on a set of representative protein folds, while DeepFold(L) is only trained on a subset of protein fold space in the cross validation.Furthermore, we study whether long-range structural motifs are useful for improving retrieval. Both FragBag and DeepFold(L) only consider local structural motif representation, while DeepFold considers both local structural motif and long-range protein contact patterns at the same time, which is harvested by the convolutional structure and non-linearity of the networks. The retrieval accuracy of DeepFold is significantly better than both DeepFold(L) and Fragbag, thus indicating that long-range structural motifs should be considered for representing protein structures.

#### Results on searching recently released proteins in PDB

In addition to the cross-validation on SCOP, we hope to further evaluate the generalization performance of DeepFold on the newly release protein structures in PDB. We download all recently released protein structures on the PDB website [1] from March 1^*st*^ 2016 to May1^*st*^ as query proteins and filter them with sequence identity 40%. Furthermore, to avoid potential redundancy between these proteins and the proteins from the SCOP database, we also remove all queries with a sequence identity higher than 40% to any protein in SCOP dataset. Then final query dataset we get has 757 proteins. By searching every query protein against all SCOP protein domains as the database, we report the retrieval performance using the same evaluations as reported above. The final DeepFold model is used by the ensemble of five models trained in the earlier 5-fold cross-validation on SCOP.

The results are reported in Figure 5 and Table 2. The curves of SGM and SSEF are not shown in the figure since their accuracies are much worse than DeepFold and FragBag.

**Figure 5:**
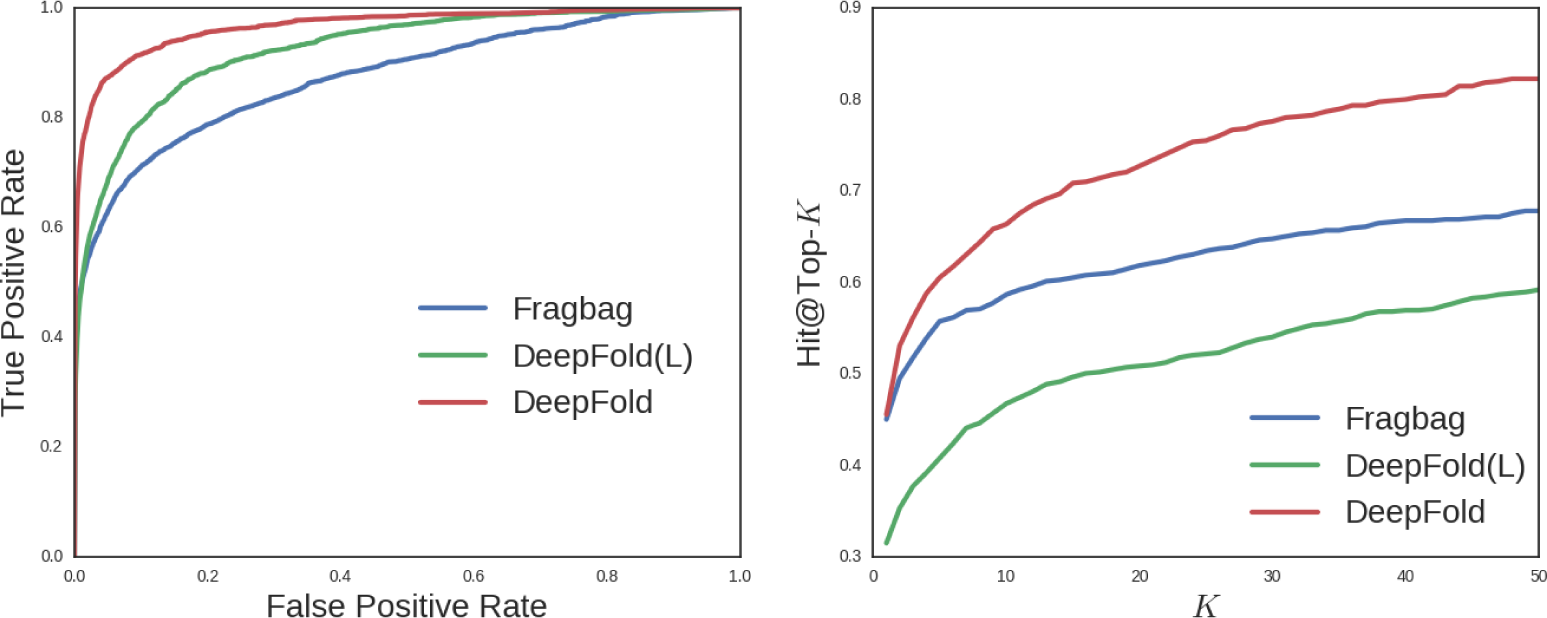
ROC curves and Hit@Top-*k* curves on the recently released proteins.

**Table 2:** Performance comparison on the recently released proteins.

| Algorithm | Dimension | AUROC | AUPRC | Top1 Acc | Top5 Acc | Top10 Acc |
| --- | --- | --- | --- | --- | --- | --- |
| SGM[14] | 30 | 0.698 | 0.048 | 0.0 | 0.0 | 0.003 |
| SSEF [31] | 1500 | 0.800 | 0.008 | 0.004 | 0.009 | 0.021 |
| FragBag [3] | 400 | 0.876 | 0.301 | 0.450 | 0.558 | 0.587 |
| DeepFold (L) | 400 | 0.925 | 0.228 | 0.315 | 0.408 | 0.468 |
| DeepFold | 400 | **0.967** | **0.375** | **0.456** | **0.605** | **0.663** |

Similar to previous results on SCOP data, DeepFold achieves better performance under all evaluation metrics on this independent non-redundant dataset, which means that the learned structural motif in our DeepFold actually can be generalized for newly released proteins that are not included in the training data. As shown in Figure 5, our Deepfold outperforms other methods by a significant margin. We also discover the wide margin between our curve and baselines in the hit@top-*k* rate. Note that though our DeepFold is only slightly better than FragBag on hit@top-1 accuracy, we are still able to obtain significant improvement on hit@top-5 and hit@top-10, which further demonstrates the effectiveness of DeepFold on protein structure search. The difference of the performance on SCOP and newly release proteins may be caused by different distributions of folds appearing in these two datasets.

#### Computational Efficiency

For computational efficiency, we evaluate the runtime needed to generate the representations of all 757 protein structures in the recently released structures curated above. On average, FragBag, one of the most efficient fingerprint algorithm before[3] took 553.202 seconds while DeepFold only took 20.128 seconds. DeepFold is roughly 25 × faster than FragBag. We argue that our algorithm is faster because it does not need to perform the expensive local structural alignments between the target protein and fragments in the library. The other reason is that DeepFold fully utilizes the GPU parallelization and the fast CUDA numerical library, even though it has many parameters in the neural network model. On CPU, DeepFold took 282.801 second, which is still faster. It is worth noting that we tried to implement Fragbag to run on a GPU, but the structure of local structural comparison algorithm makes it tough to fully exploit the parallelism of GPU cores to outperform the implementation running on CPU.

### 4.2 Why does DeepFold work so well?

To interpret the superior performance DeepFold achieves in protein structure search, we analyze the representation learned by the deep neural network model and intend to understand the structural meaning of the representation. To do so, we develop a quantitative approach to visualize the structural motifs learned in this deep neural network. In detail, we firstly obtain the activation values of filters in each convolutional layer for each protein in a non-redundant protein structure dataset. Then for each filter, we obtain the top proteins with the highest activation values and extract the corresponding protein substructures. These substructures are aligned together, and a central substructure is picked to represent the motif learned by the filter.

To visualize structural motifs in a well-organized way, we utilize t-SNE[11] to perform a nonlinear dimension reduction, projecting these motifs into a 2D space. Figure 6 illustrates the structural motifs we learned in DeepFold, where each point denotes a learned structural motif. The left plot presents all local contiguous structural motifs, while the right one presents long-range interacting motifs. We annotate each point with a specific color indicating the secondary structure composition of the corresponding motif, where blue indicates *α*-helix, red indicates *β*-strand and the purple one indicates loop. From the left plot, we could observe the local structural motifs include not only *α*-helix, *β*-strand, but also some mixture of short *α/β* structures. In addition, the long-range structural motifs are mainly *β*-*β* interacting patterns, under the complicated environment with *β*-strand, *α*-helix or even loop structure. Furthermore, we observe that the low-level motifs learned by DeepFold can be organized to represent high-level structural motifs by composing local and long-range motifs together. From this visualization, it is clear that DeepFold extracts not only meaningful contiguous backbone fragments but also the long-range interacting motifs, which are entirely ignored in FragBag. It is worth noting that FragBag can be seen as a special case of DeepFold if we only consider the near-diagonal region of the pairwise distance matrix as input.

**Figure 6:**
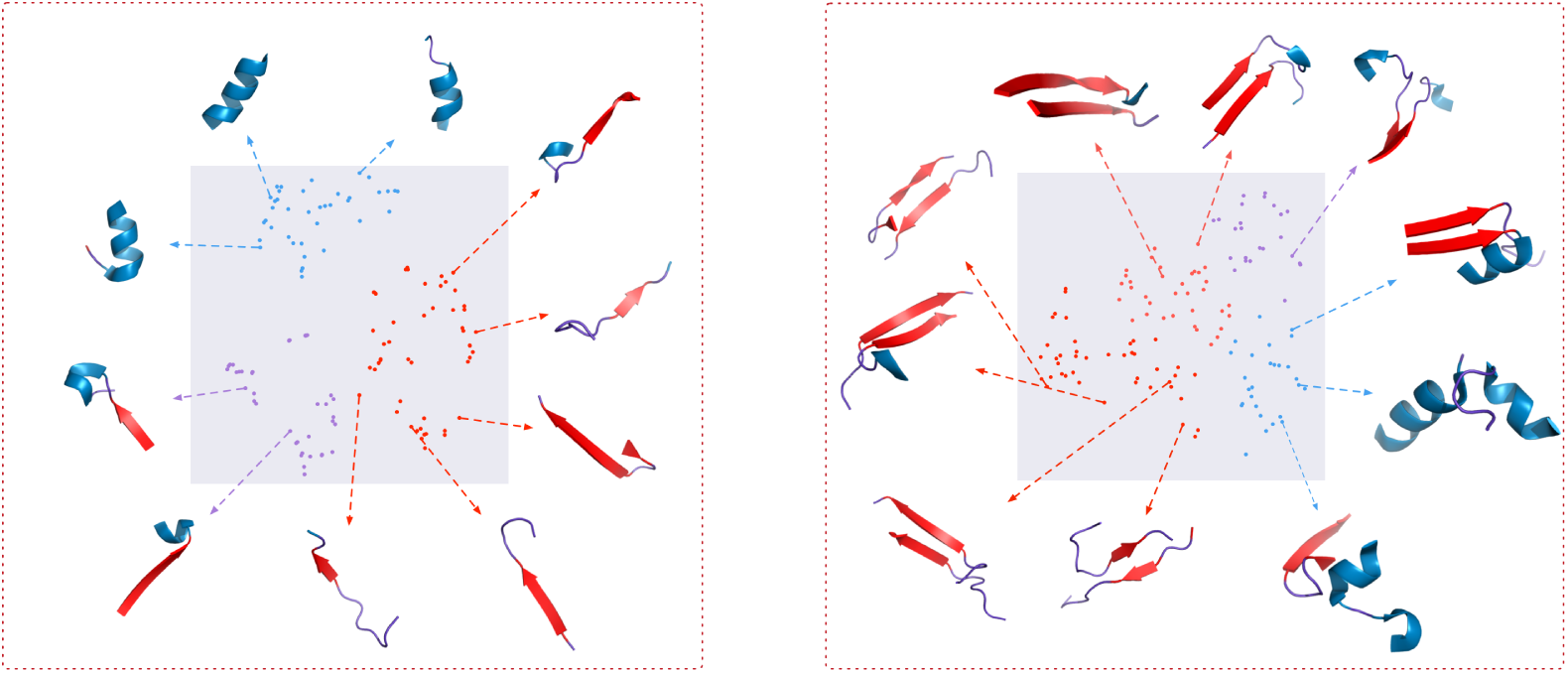
Structural motifs learned by DeepFold. The left figure shows the local backbone-segment motifs, and the right figure shows the long-range interacting contact motifs. Motifs are clustered and visualized according to their secondary structural composition. 10 representative motifs are shown with their structures.

To compare the structural motifs against the fragment library used by FragBag. We visualize the carbon backbones of the motifs and fragments in Figure 8. We observe that the local structural motifs learned by DeepFold are similar to the fragments utilized by FragBag. Also, the DeepFold preserves long-range contacts in the original structures, which contribute to a more powerful structural representation.

In addition to visualizing the structural motifs learned by DeepFold, we also check what are the “false-positive” hits found by DeepFold. From the Figure 7, we observe that the “wrongly” predicted top structures also share substantial structural similarity with the query proteins, indicating that our DeepFold is potentially capable of finding remotely related structural neighbors which contains similar motif composition.

**Figure 7:**
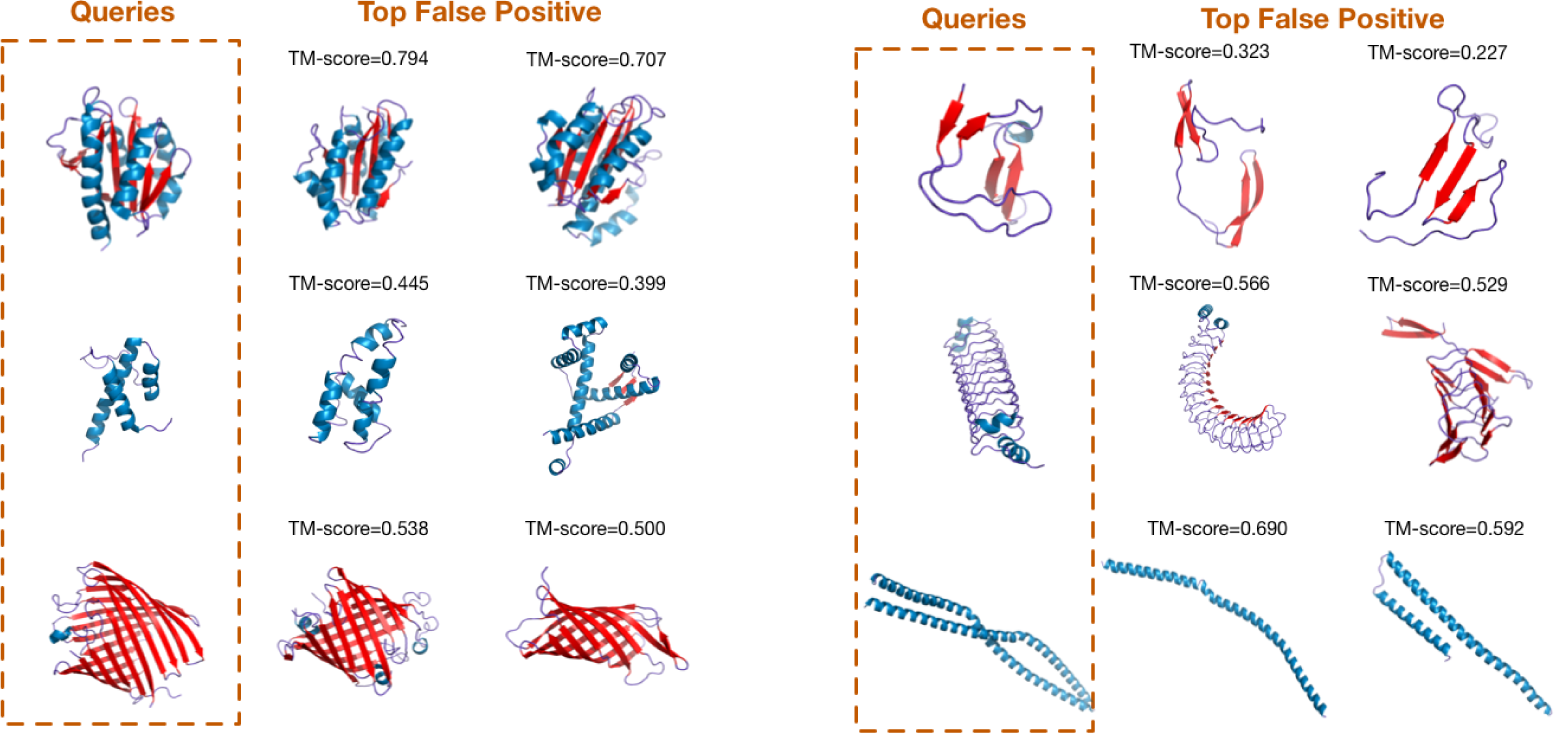
“False positives” predicted by DeepFold.

**Figure 8:**
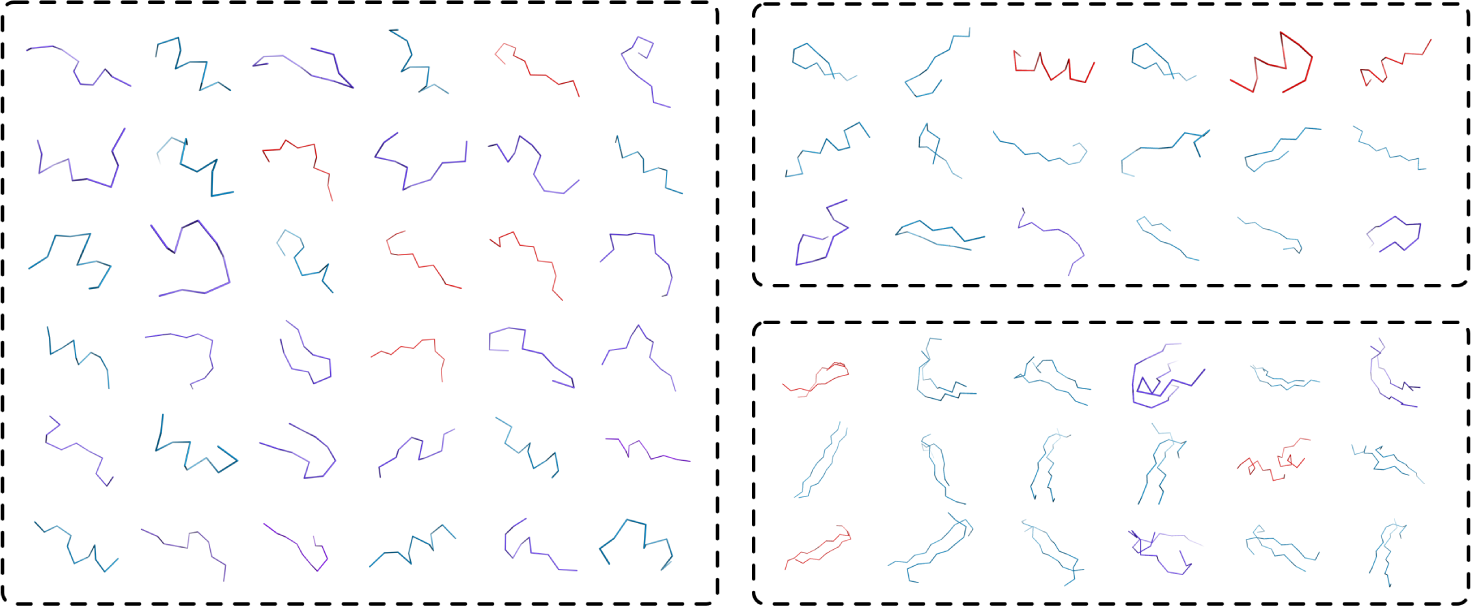
Comparison of motifs used by DeepFold and FragBag. The left panel shows fragments used by FragBag. The upper right panel shows local structural motifs, and the bottom right shows long-range contact motifs by DeepFold.

## 5 Conclusion

In this paper, we present DeepFold, a deep-learning approach to building structural motif presentations for better alignment-free protein structure search. We develop DeepFold, a deep convolutional neural network model to extract effective structural patterns of a protein structure from its *C*_*α*_ trace. Similar to the previous FragBag, we represent each protein structure/fold using a vector of the learned motifs and perform the structure search by only computing vector similarity. This neural network is trained in a supervised manner by discriminating similar template structures from dissimilar structures in a database. We demonstrate that DeepFold greatly outperforms FragBag on protein structure search on SCOP database and a set of newly released PDB structures, regarding both search accuracy and efficiency for computing structural representations. Remarkably, after visualizing the motifs learned by DeepFold, we find that it not only extracts meaningful backbone segments but also identifies important long-range interacting structural motifs for structural comparison. Furthermore, given that the retrieval accuracy of DeepFold is outstanding, we can combine it with structural alignment algorithms in the compressive genomics manner by first applying DeepFold as a “coarse search” step to identify a very small subset of putative similar template structures that are ranked very top, and then performing structural alignment algorithms as a “fine search” step to refine the ranking and obtain more informative residue-level structural alignments for downstream analysis [25]. Finally, we expect that the structural motifs extracted by DeepFold will provide new insights into the evolution and hierarchical organization of protein structure.

## Footnotes

1 https://github.com/Lasagne/Lasagne

